# Spatial proteomics reveals mechanisms of cell-intrinsic tryptophan metabolism controlling ovarian cancer survival

**DOI:** 10.1101/2025.08.14.670277

**Authors:** Teng Teng Xu, Andreas Metousis, Lisa Kainacher, Xin Zhang, Barbara Steigenberger, Katherine G. Madden, Lisa Schweizer, Charlotte Duteil, Sofia Rossini, Ernst Lengyel, Eva Obermayr, Ziv Shulman, Thierry M. Nordmann, Eric L. Lindberg, Matthias Mann, Peter J. Murray

## Abstract

Indole-2,3-dioxygenase (IDO1) depletes tryptophan to dampen anti-tumor T cells, yet IDO1 inhibitors (IDO1i) have failed clinically. Using deep visual proteomics, we isolated IDO1 high, medium and low ovarian tumor cells in situ and found IDO1 tightly linked to interferon-γ (IFN-γ) signaling and heterogeneously expressed. Across orthogonal models with tunable IDO1, IFN-γ killed ovarian cancer via a pathway requiring IFN-γ signaling, IDO1-dependent tryptophan depletion, and a biphasic integrated stress response that initially protects from starvation and later drives death. IDO1i or tryptophan supplementation rescued these effects, promoting tumor survival. These data reveal a context-dependent, tumor-suppressive facet of IDO1 and explain how IDO1i can paradoxically favor cancer viability. Our findings call for re-evaluation of IDO1 as a target and suggest exploiting the tryptophan-starvation/GCN2-ISR axis to enhance therapy.

## Main text

IDO1 and TDO2 are heme enzymes that degrade tryptophan (TRP) by oxidation of the indole ring, initiating the kynurenine pathway (*1, 2*). In normal animals, TDO2 is constitutively expressed in the liver and regulated by the ubiquitin-proteasome system (*3*). By contrast, IDO1 expression is inducible by interferons (IFNs) and especially IFN-γ, which links its expression and function to inflammation (*4*). A key function of IDO1 is to create a TRP depleted environment to block virus and intracellular parasite replication (*1, 5, 6*). IDO1-mediated tryptophan depletion is also tied to fetal tolerance in pregnancy and blockade of T cell proliferation (*1, 7, 8*). The latter pathway was the trigger for the development of IDO1 inhibitors against cancer (*9, 10*). In this context, IDO1-expressing tumor and/or immune cells within the tumor microenvironment (TME) are proposed to deplete local TRP (*11*). The resulting local TRP deficiency leads to impaired anti-tumor T cell proliferation and function, thereby aiding tumor immune evasion. (*9-11*). Thus, an IDO1i should restore local TRP and promote anti-tumor T cell immunity. However, despite strong preclinical rationale, all IDO1i have failed in clinical trials (*12, 13*).

Recently, it was uncovered that IDO1 activates additional pathways implicated in cancer survival or progression. These pathways include de novo NAD^+^ biosynthesis and maintenance of genome stability (*14*), 1-carbon metabolism (*15*), activation of the aryl hydrocarbon receptor (AHR) by IDO1 metabolites (*16*) and ferroptosis suppression via a pathway that involves kynurenine products controlling cystine importation and free radical scavenging (*17*). Furthermore, IDO1^+^ melanoma cells were recently shown to be sensitive to attack by engineered antigen-specific T cells and IFN-γ production (*18*). Notably, IDO1i reversed the lethal effects of the T cells via a mechanism involving IFN-γ and IDO1-dependent expression of the MITF transcription factor. While that study provided an important clue as to why IDO1i failed in clinical trials and could have unwanted effects opposite to the original target validation approach, we sought to understand the role of IDO1 and the consequences of its inhibition in cancer by quantifying the proteomes of tumor cells expressing IDO1 directly in their native microenvironment.

To investigate this IDO1 paradox, we focused on ovarian cancer, a tumor type where few therapeutic options are available and IDO1 is highly and heterogeneously expressed across different cells within the TME. (*19*). We hypothesized that tumor cell-intrinsic IDO1 is a critical factor that controls tumor cell viability, distinct from effects on T cells or other cells in the TME. To test this, we required a methodology that could resolve IDO1 expression at the single-cell protein level while preserving spatial context. Conventional bulk tissue analysis would average IDO1^hi^ and IDO1^lo^ cells, masking specific cell-intrinsic effects. We therefore, employed deep visual proteomics (DVP), which allowed us to isolate and analyze individual tumor cells based on their IDO1 expression levels directly from patient tissues, thereby capturing, rather than diluting, the true heterogeneity of IDO1 expression.

## Results

### DVP workflow of high grade serous ovarian cancer

High grade serous ovarian cancer (HGSOC) is a tumor frequently associated with IDO1 expression (*19*) in which IDO1i (epacadostat) failed to improve clinical outcomes (*20*). To investigate the characteristics of IDO1^+^ ovarian tumor cells, we used DVP (*21, 22*). We collected treatment-naive HGSOC samples from four individuals and stained formalin-fixed, paraffin-embedded (FFPE) sections with specific antibodies to IDO1, EPCAM (for tumor cells) and CD45 (all hematopoietic cells). Deep learning-based segmentation of cells within the FFPE sections allowed isolation of EPCAM^+^ cell shapes while excluding the CD45^+^ immune cells. Within the EPCAM^+^ tumor cells, we applied k-means clustering to differentiate between IDO1^hi^ and IDO1^lo^ cell shapes and additionally isolated IDO1^+^ cells (IDO1^med^) with intermediate signal (Fig. 1A). This approach allowed us to compare the proteome of tumor cells across IDO1 expression, from high via mid to low, within the same tumor sample.

**Fig. 1.**
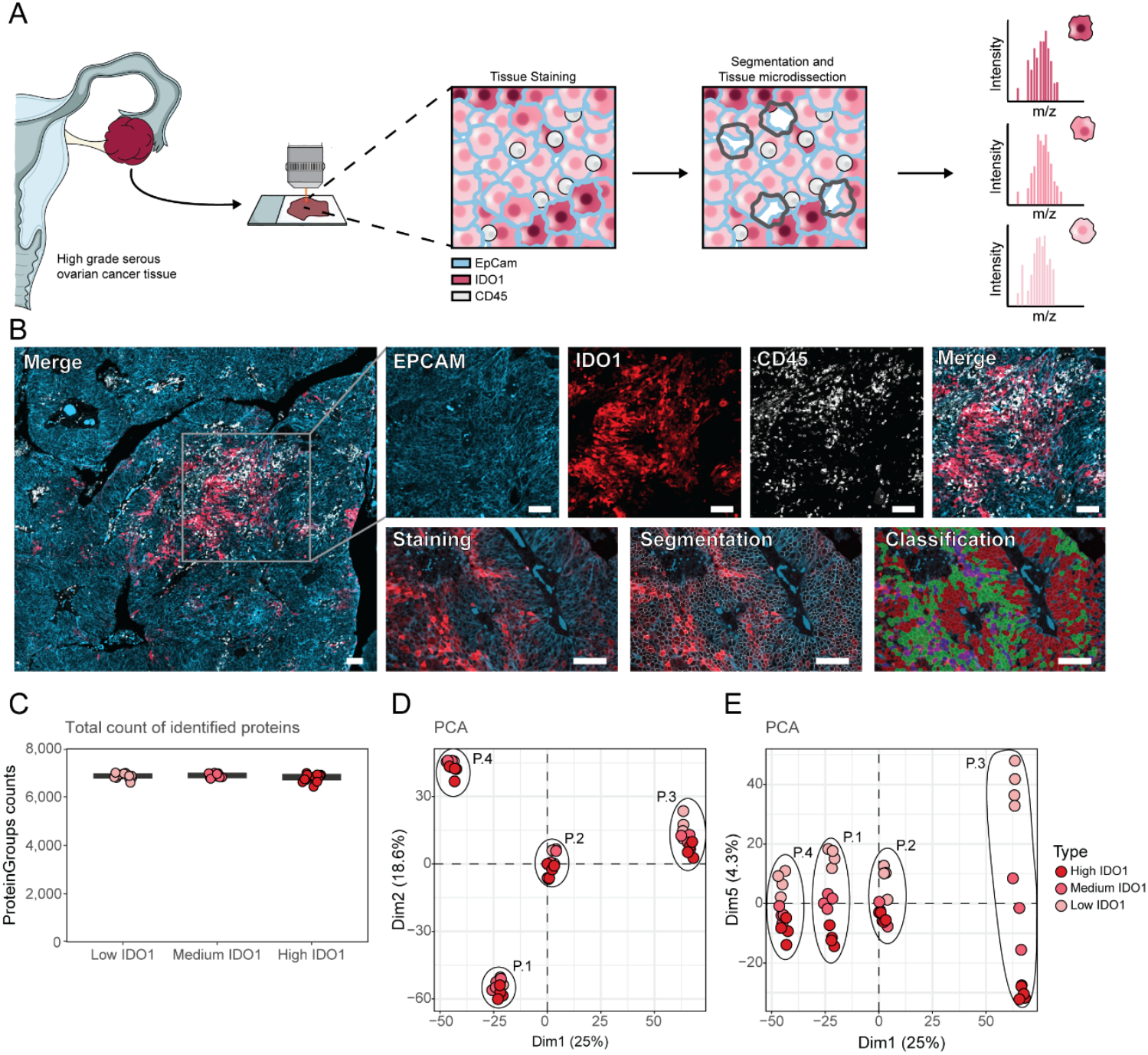
Deep visual proteomics workflow reveals IDO1-driven cancer cell heterogeneity. (**A**) DVP workflow schematic. HGSOC tissues stained for IDO1, EPCAM (tumor cells), and CD45 (immune cells), followed by cell segmentation and k-means clustering to isolate IDO1^hi^, IDO1^med^ and IDO1^lo^ tumor cells for deep proteomics analysis. (**B**) Representative immunofluorescence images showing heterogeneous IDO1 expression in patient 1. Bottom panels show computational segmentation and classification workflow. Scale bars = 100 μm. Cyan: EPCAM, red: IDO1, white: CD45. In the classification image (bottom right) red color corresponds to cells classified as IDO1^lo^, green to IDO1^med^ and blue to IDO1^hi^. (**C**) Total protein identification counts across IDO1 expression groups. (**D**) PCA of proteome data. PC1 and PC2 capture patient heterogeneity (P1-P4). (**E**) PCA showing IDO1-driven expression patterns. PC5 separates samples by IDO1 levels: red (high), pink (medium), light pink (low).

### Proteome of IDO1^hi^ and IDO1^lo^ ovarian cancer cells

The spatial distribution of IDO1^+^ tumor cells in each sample, as visualized by immunofluorescence, was heterogeneous. Clusters of IDO1^hi^ cells adjacent to IDO1^med^ and IDO1^lo-^ cells were scattered throughout the tumor bed in all samples; a property of HGSOC previously uncovered by single-cell RNA-seq approaches as discussed below (Fig. 1B and S1A). Thus, DVP in this tumor context allowed us to examine the proteomes in a cell type resolved level, without disrupting the native architecture of the TME. We identified ∼6500 proteins in each analyzed group, each of which contained ∼100 cell equivalents, with high data completeness and and high reproducibility (Fig. 1C, S1B-C). Principal component analysis (PCA) captured patient heterogeneity in principal component 1 (PC1), explaining 25% of the sample variability. (Fig. 1D), while PC5 revealed a clear separation driven by IDO1 status (Fig. 1E). In every case, IDO1 was the most differentially identified protein in the IDO1^hi^ versus IDO1^lo^ comparison, validating the fidelity of the methodology (Fig. 2A and Fig. S1D-G).

**Fig. 2.**
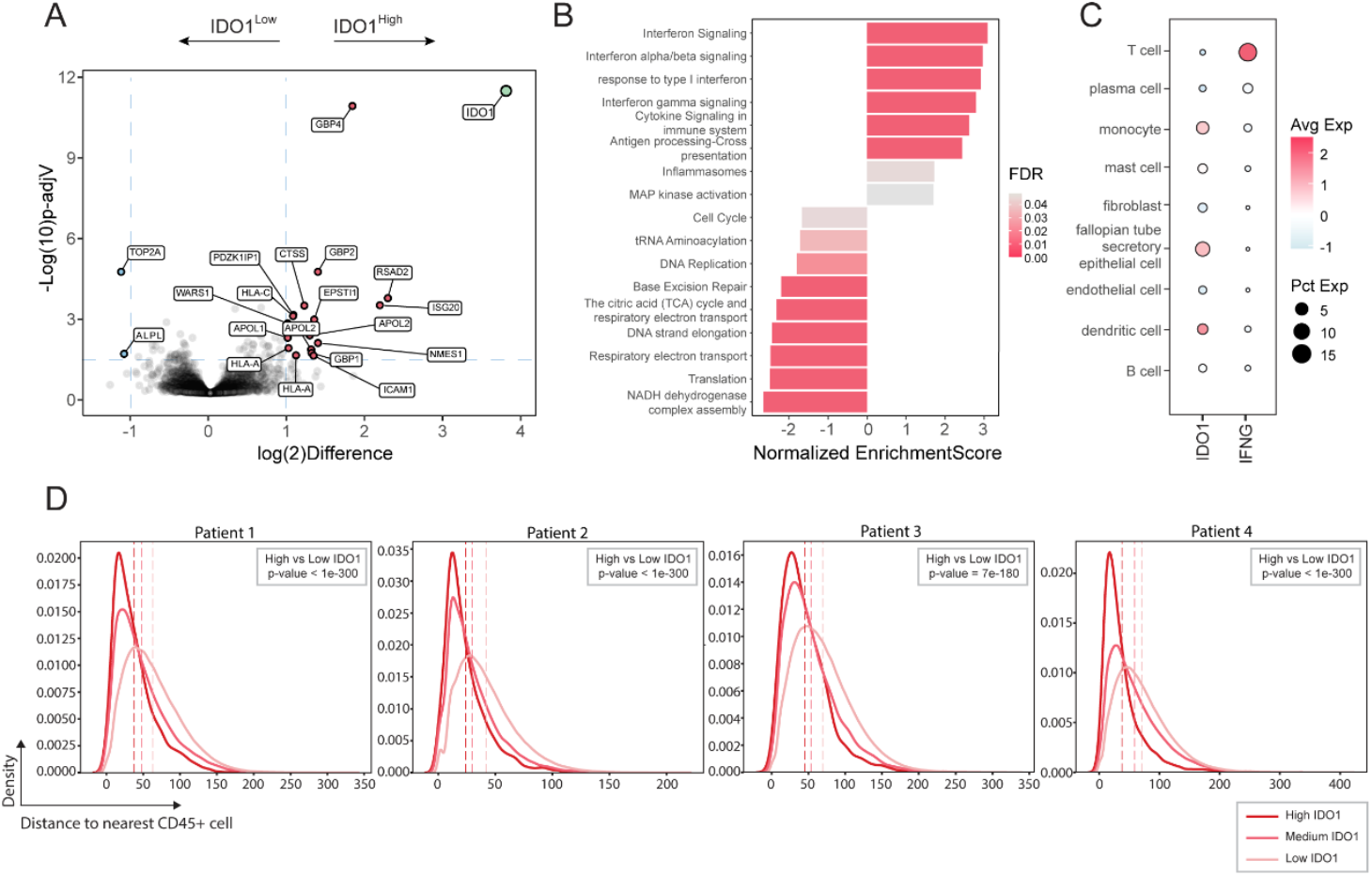
IDO1^hi^ tumor cells exhibit IFN-γ response signatures and spatial proximity to immune cells. (**A**) Volcano plot showing differentially expressed proteins between IDO1^hi^ and IDO1^lo^ tumor cells (pooled data from 4 patients). Gray dots represent non-significant changes; thresholds for significance are log(2)Difference < -1 or >1 and - Log(10)p-adjV > 1.3. (**B**) Gene set enrichment analysis (GSEA) revealing pathways of differentially expressed proteins in IDO1^hi^ versus IDO1^lo^ tumor cells. Bar length represents normalized enrichment score; color intensity indicates enrichment significance. Positive values indicate enrichment in IDO1^hi^ cells. (**C**) Dotplot shows scaled and averaged expression levels of IDO1 and IFNG mRNAs per cell type in the TME. Dot size reflects the fraction (%) of expressing cells; color scaled means expression level. (**D**) Spatial proximity analysis between tumor cells and CD45^+^ immune cells across patients P1-P4. Density plots show the distribution of distances (in arbitrary BIAS units) from each tumor cell to the nearest immune cell, stratified by IDO1 expression: red (high), pink (medium), light pink (low). Dashed lines represent mean distances per group. Statistics from two-sample t-tests comparing IDO1^hi^ versus IDO1^lo^ groups are shown in the top right corner of each plot.

We next asked which pathways were associated with IDO1. Gene set enrichment analysis (GSEA) across all patients revealed that nearly all the proteins that were associated with the IDO1^hi^ group (e.g. GBP4, GBP2, RSAD2, HLA-C, HLA-A, ICAM-1 etc.) belonged to the IFN-γ-response and inflammatory pathways (Fig 2A and Fig. 2B), while showing no evidence to support an AHR signature (*23-25*). On the other hand, These data suggest that the clusters of IDO1^+^ cells in each HGSOC samples had been stimulated by IFN-γ. Previous studies in melanoma reached a similar conclusion; IDO1 expression is tightly linked to IFN-γ, most likely in localized niches within the TME (*26, 27*). On the other hand, pathways linked to DNA replication, translation and cell cycle were enriched in the IDO1^lo^ group (Fig. 2B), suggesting that IDO1^lo^ tumor cells might proliferate more than the IDO1^hi^. At first, such a proliferative proteome profile of IDO1^lo^ tumor cells might appear surprising, as IDO1 expression has generally been associated with poor prognosis in cancer (*9, 28-30*). However, our findings are in line with recent reports, where IDO1 seems to be anti-tumorigenic (*18, 31*).

### Single cell RNA sequencing analysis of IDO1 positive tumor cells

We next investigated the relationship between IDO1 and IFN-γ expression in HGSOC using a large single-cell RNA sequencing (scRNA-seq) dataset from 42 treatment naive patients for a total of 160 biopsies (*32*). In this dataset, we found a strong correlation between IDO1 expression and IFN-γ response, reflecting our spatial proteomics data (Fig. S2A). Moreover, IDO1 expression was detected in tumor and myeloid cells (Fig S2B and Fig. S2C). As our focus was on tumor cell IDO1 expression and function, we next asked which cells expressed IFN-γ in the same dataset. Consistent with previous studies in TMEs, T cells were the predominant source of IFN-γ mRNA (Fig. 2C). HGSOC cells in our study and in the scRNAseq database show high heterogeneity in IDO1 expression not only across different patients (Fig.S2D), but also within the same tissue (Fig. 1B and Fig. S1A). These data support a model in which T-cell derived IFN-γ induces IDO1 expression in spatially proximate, responsive cells (*33*), rather than a cell-intrinsic pathway (*34*). To test in our cohort if there is a relationship between the proximity of tumor cells relative to the proposed cellular source of IFN-γ and IDO1 expression, we measured the distance of the different IDO1 expressing tumor cells from the nearest immune cell (CD45^+^). We found a strong inverse correlation between IDO1 expression levels and distance from the immune cells in all our analyzed patients (Fig. 2D).

### IFN-γ induces IDO1 dependent starvation via the GCN2 pathway

To better understand the role of IDO1 in ovarian cancer we needed an experimental platform where we could tune IDO1 expression. We therefore screened a panel of validated human ovarian cancer lines for IDO1 expression. We selected three groups of ovarian cancer cells: (i) lines with constitutive IDO1 expression where IDO1 levels can be further increased by IFN-γ (SKOV3, OVCAR4), (ii) lines devoid of baseline IDO1 expression but with high IFN-γ-inducibility (OVCAR3, OVCAR5) and lines that lack IDO1 expression at baseline and upon IFN-γ treatment (OVCAR8, PA-1) (Fig. 3A).

**Fig. 3.**
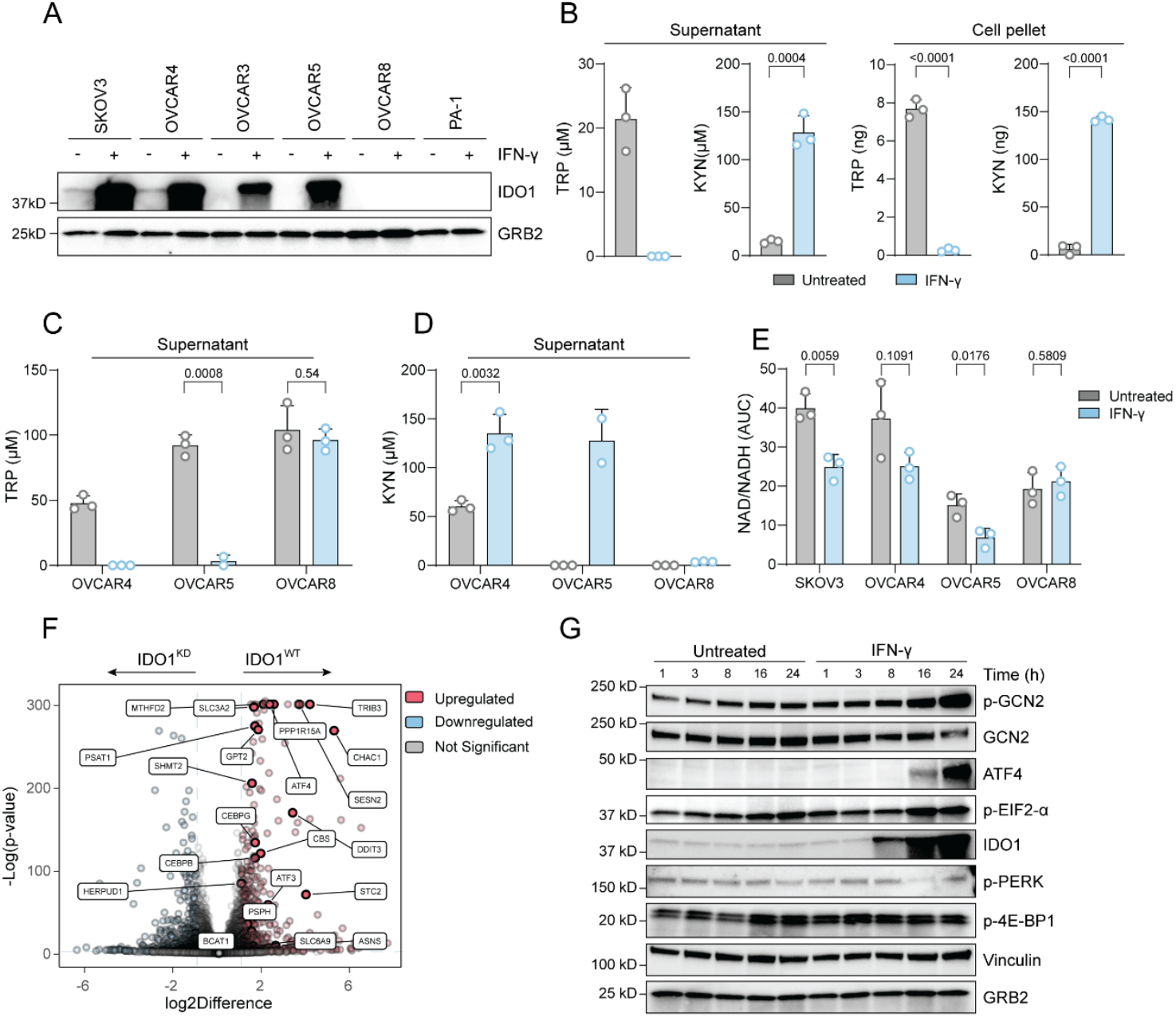
IDO1 converts TRP to KYN and activates the ISR. (**A**) Immunoblot analysis for IDO1 of different ovarian cancer cell lines +/-10 ng/ml IFN-γ. (**B** to **D**) TRP and KYN levels measured by mass spectrometry in the supernatant or in the cell pellet of SKOV3 (B), TRP in the supernatant of OVCAR4, OVCAR5 and OVCAR8 (C) and KYN in the supernatant of OVCAR4, OVCAR5 and OVCAR8 (D). All cells were stimulated with 10 ng/ml IFN-γ for 24hr. (**E**) NAD and NADH ratio measured by mass spectrometry in the cell pellets of SKOV3, OVCAR4, OVCAR5 and OVCAR8 stimulated with 10 ng/ml IFN-γ for 24hr. (**F**) Volcano plots showing overall changes in the transcriptome in SKOV3^WT^ versus SKOV3 IDO^KD^ both stimulated with 10 ng/ml of IFN-γ for 24hr. Annotated genes belong to the ISR pathway. Thresholds for significance are log(2)Difference <-1 or >1 and -log(p-value) > 2. (**G**) Time-course immunoblot of SKOV3 +/-10 ng/ml IFN-γ. All data are shown as means ± SD of three independent experiments, exception for (C) and (D) with only two replicates in OVCAR5. The unpaired t-test was used in (B), (C), (D) and (E), statistics was not performed in conditions where the metabolites were below the detection limit and values were arbitrary set to 0.

IDO1 oxidizes TRP to N-formyl-kynurenine to initiate the kynurenine (KYN) pathway (KP), which eventually leads to de novo NAD^+^ biosynthesis and production of TCA cycle and 1-carbon intermediates (*2, 15, 35, 36*) (Fig. S3A). Using mass spectrometry to measure the activity of these pathways in ovarian cancer cells treated with IFN-γ, we found TRP depletion and KYN production correlated with IDO1 expression; OVCAR8, which does not express IDO1 also did not produce detectable KYN. SKOV3, OVCAR4, OVCAR5 all depleted TRP to undetectable amounts following IFN-γ treatment within 24 hr (Fig. 3B-D and Fig. S3B-C).

We next investigated the relation between IFN-γ and two pathways that could plausibly control IDO1-dependent outcomes in ovarian cancer: de novo NAD^+^ biosynthesis and ferroptosis suppression. First, IFN-γ reduced NAD^+^/NADH ratio rather than increasing it (Fig. 3E). Further, FK866, a NAMPT inhibitor, which blocks the NAD^+^ salvage pathway from nicotinamide (*14*), killed ovarian cancer cells, suggesting that ovarian cancer cells rely more on the salvage pathway as source for NAD^+^ rather than the IDO1-mediated de novo biosynthesis pathway (Fig. S3D). Next, we assessed the relationship between KP pathway metabolites and ferroptosis protection. Our previous work proposed that IDO1^+^ cells increase KP products locally and protects the nearby cells from redox stress and ferroptotic cell death (*17*). Thus, IDO1^KD^ cells should be more sensitive to ferroptosis inducers than cells that can generate KYN. We observed a poor protection of both SKOV3 and OVCAR4 by KYN, on the contrary 3-hydroxy-kynurenine (3-HK) and hydroxy-anthranilic acid (HAA), downstream metabolites of the KP, were strong inhibitors of ferroptosis (Fig. S3E and S3G). Despite these results, data from the SKOV3 and the OVCAR4 were inconsistent, the first showed no dependence to IDO1 in terms of sensitivity to ferroptosis inducers (FINs), while the latter displayed some difference (Fig S3F and S3I). We also measured the key anti-ferroptosis KP metabolites, 3-HK and HAA, which were below the limit of detection. Therefore, we had insufficient evidence supporting a role for IDO1 in either de novo NAD^+^ biosynthesis or ferroptosis suppression in ovarian cancer. Notably, our experiments on ferroptosis resistance or sensitivity were conducted on ovarian cells in the absence of IFN-γ stimulation, as we had coincidently observed that IFN-γ triggered cell death. We therefore investigated if IFN-γ and IDO1 conspired to control ovarian cancer cell viability.

To investigate alternate cellular pathways controlled by IDO1 relevant to ovarian cancer, we generated polyclonal SKOV3, OVCAR4 and OVCAR5 cells depleted of IDO1 using CRISPR/Cas9. We treated control SKOV3 or SKOV3-IDO1 knockdown (IDO1^KD^) cells with IFN-γ and compared them by RNA-seq, from which a striking pattern emerged. Transcripts induced in the WT SKOV3 cells compared to the IDO1^KD^ cells were almost exclusively associated with the GCN2-dependent amino acid stress response (Fig. 3F). Kinetically, IDO1 expression was apparent 8 hr after IFN-γ stimulation and was followed by an increase of GCN2 phosphorylation 8 hr later (p-T899, the key activation phosphorylation event for the GCN2 pathway) (*37*) and expression of the downstream pathway from the GCN2 integrated stress response (ISR), including phosphorylation of EIF2-α and ATF4, the transcription factor that controls the majority of the ISR (*38*) (Fig. 3G). Activation of the ISR is a protective pathway that reduces translation and activates transcripts and proteins that are cell protective (*38*). However, when we treated SKOV3 cells with IFN-γ to induce IDO1 expression, we discovered that the IFN-γ-treated cells slowly died, beginning at around 4-5 days (Fig. 4A, 4B and 4C). This observation prompted us to investigate whether IFN-γ–induced IDO1 controls time-dependent tumor cell survival and whether it is linked to the GCN2-dependent starvation pathway.

**Fig. 4.**
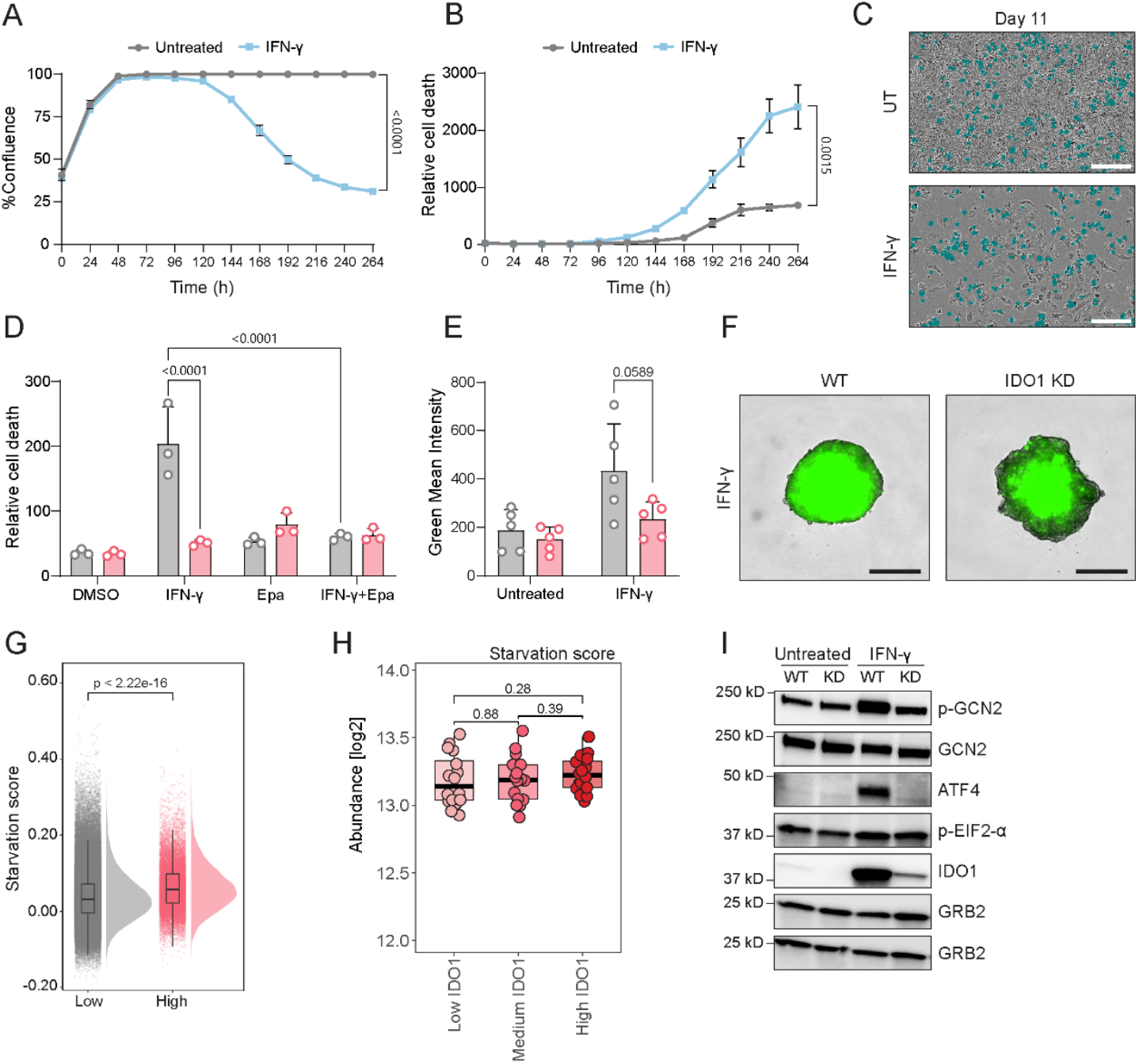
Prolonged IFN-γ stimulation promotes IDO1 dependent ISR induction and cell death. (**A** and **B**) Time-course live cell imaging monitoring cell confluence (A) and relative cell death (CellTox Green positive cells normalized to confluence) (B) of SKOV3 treated with 10 ng/ml IFN-γ. (**C**) Representative bright-field image of SKOV3 +/-10ng/ml IFN-γ at day 11. Scale bar = 200 μm (**D**) Quantification of live cell imaging data as relative cell death of SKOV3^WT^ or SKOV3 IDO^KD^ treated for 5 days with either vehicle (DMSO), 10 ng/ml IFN-γ, 10 µM epacadostat (Epa) or the combination of the two. (**E**) Green fluorescence intensity measured with live cell imaging of SKOV3^WT^ or SKOV3 IDO^KD^ spheroids cultured for 5 days +/-10 ng/ml IFN-γ. (**F**) Representative images of (E) at day 5. Scale bar = 300 μm (**G**) Density curves and boxplots show starvation score per tumor cell in the IDO1^lo^ and IDO1^hi^ expressing group. Significant p-value was calculated with the Wilcoxon rank-sum test. (**H**) Starvation pathway activation in patient-derived IDO1^hi^, IDO1^med^, and IDO1^lo^ tumor cells (pooled data from 4 patients). Starvation scores calculated by averaging log2 abundance levels of starvation-associated proteins. Statistical comparisons performed using paired two-sample t-tests with equal variances. **(I)** Immunoblot analysis of the ISR pathway in SKOV3^WT^ versus SKOV3 IDO^KD^ untreated or stimulated with 10 ng/ml IFN-γ for 24hr (n = 3). Representative plot of three independent experiments, data are shown as means ± SD of three technical replicates and statistics was calculated using the unpaired t-test comparing data at day 11 in (A) and (B). Data are shown as means ± SD of at least three independent experiments and two-way Anova complemented with a Tukey’s multiple comparison was used in (D) and (E).

### IFN-γ killing is regulated by IDO1-dependent TRP starvation

The first question we addressed was whether IDO1 was required for the IFN-γ-mediated killing of SKOV3 cells. We conducted a cell death assay comparing SKOV3^WT^ and IDO1^KD^ exposed to IFN-γ for 5 days. As an additional control we included epacadostat, a key IDO1 inhibitor that has been extensively used in clinical trials (*13*). Indeed, IDO1 depletion or inhibition protected SKOV3 cells from IFN-γ killing (Fig. 4D and Fig. S4A). Both OVCAR4 and OVCAR5 were also sensitive to IFN-γ, while their IDO1^KD^ counterparts were more resistant (Fig. S4B and S4D). OVCAR8 and PA-1, IDO1 negative cells, were resistant to IFN-γ-induced cell death (Fig. S4E and S4F). The same pattern was observed when SKOV3 cells were cultured in 3D spheres to mimic better the complex tumor architecture *in-vitro* (Fig 4E and 4F). Further, we evaluated the HGSOC patient DVP and scRNAseq data to obtain proxies for the activation of the GCN2 pathway (Table S1). We observed a statistically significant increase of the GCN2 pathway at the RNA level, consistent with fact that the ISR activates transcription of hundreds of mRNAs (Fig. 4G). However, as the ISR coincidently inhibits global translation with the exception of a small number of key induced targets, the proteomes of IDO1^hi^ tumor cells did not shown differences compared to IDO1^lo^ cells (Fig. 4H).

Next, we compared the p-GCN2 response in IDO1^WT^ and IDO1^KD^ cells in response to IFN-γ and found it was IDO1 dependent. (Fig. 4I and S4C). We reasoned that if TRP depletion is the driving cause of cell death, TRP supplementation should rescue the cells. Doubling the amount of the normal media TRP prevented cell death and GCN2 phosphorylation, consistent with our targeted metabolomics data showing complete conversion within a few hours of TRP to KYN once IDO1 is induced (Fig. 5A and Fig. 5B). Furthermore, to uncouple the effects of TRP depletion and KYN accumulation resulting from the activity of IDO1, we cultured IDO1^KD^ cells in DMEM without TRP and measured an increase of cell death comparable to the WT cells. Meanwhile, addition of KYN to the culture medium did not affect cell viability (Fig. 5C and Fig. S4G). Hence, we ruled out a KYN-dependent effect of modulating IFN-γ-mediated death, contrasting findings from other studies where KYN was toxic to some cancer cells (*39*). We complemented these findings with the use of a JAK1/2 inhibitor (ruxolitinib) and a doxycycline-inducible IDO1 overexpression system. Ruxolitinib blocks IFN-γ signaling by inhibiting the JAKs required for its downstream signaling, including IDO1 expression. As predicted, ruxolitinib prevented IFN-γ killing (Fig.5D). Overexpression of IDO1 but not its enzyme dead version F226A (*40*) could increase cell death of the IDO1^KD^ SKOV3 cells to comparable levels as in the WT cells stimulated with IFN-γ (Fig. 5E-F and S4H). Collectively, our results suggest that IDO1 expression, induced by IFN-γ via JAK1/2 signaling, rapidly consumes TRP from the environment and triggers the p-GCN2 driven ISR, which ultimately leads to cell death.

**Fig. 5.**
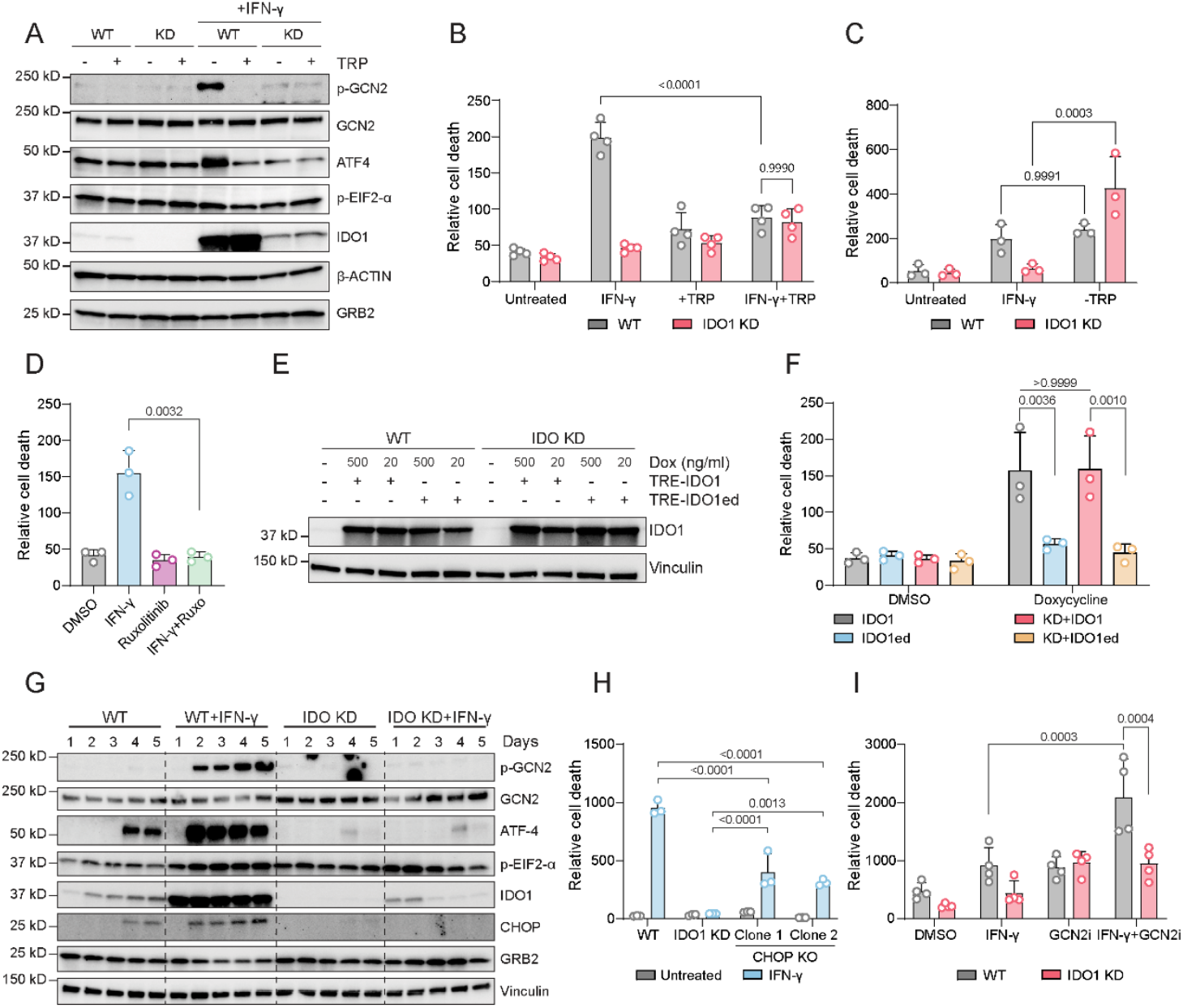
TRP depletion cell death is regulated by CHOP and GCN2 inhibition boosts tumor killing. (**A**) Immunoblot analysis of the ISR in SKOV3^WT^ versus SKOV3 IDO1^KD^ stimulated with 10 ng/ml IFN-γ for 24 hr with 156 µM TRP supplement every 12 hr (n = 3). (**B** to **D**) Quantification of live cell imaging data as relative cell death at day 5 of SKOV3^WT^ versus SKOV3 IDO1^KD^ stimulated with 10 ng/ml IFN-γ and 156 µM TRP supplement every 12 hr (B), relative cell death of the same cells as in (B) cultivated in DMEM without TRP (C) and relative cell death of SKOV3^WT^ treated with either 10 ng/ml IFN-γ, 5 µM ruxolitinib or both (D). (**E** to **F**) SKOV3^WT^ and SKOV3 IDO1^KD^ expressing either TRE-hIDO1 or TRE-hIDO1ed (F226A) stimulated with doxycycline. Immunoblot analysis for IDO1 testing 500 ng/ml and 20 ng/ml of doxycycline (E, n = 1) and relative cell death after 5 days of 20 ng/ml doxycycline stimulation every 3 days (F). (**G**) Time-course immunoblot analysis of ISR including CHOP in SKOV3^WT^ versus SKOV3 IDO1^KD^ stimulated with 10 ng/ml IFN-γ (n = 3). (**H** and **I**) Quantification of live cell imaging data as relative cell death at day 6 of SKOV3^WT^, IDO1^KD^ or CHOP^KD^ +/-10 ng/ml IFN-γ (H) and day 8 of SKOV3^WT^ and IDO1^KD^ treated with either vehicle (DMSO), 10 ng/ml IFN-γ, 1 µM GCN2-IN-6 (GCN2i) or both (I). All data are shown as means ± SD of at least three independent experiments, except for (H) where a representative plot of three independent experiments is presented, data are shown as means ± SD of three technical replicates. Two-way Anova complemented with a Tukey’s multiple comparison was used in (B), (C), (F), (H) and (I). Unpaired t-test was used in (D).

### TRP starvation-induced cell death is partially regulated by CHOP

Depletion of amino acids activates the ISR and induce cell death through CHOP (encoded by *DDIT3*) (*41-43*).To verify if this is also true for TRP starvation upon IFN-γ stimulation, we monitored ISR activation, including CHOP expression, over the full time-frame of our cell death assay. We detected a stable ISR activation and CHOP upregulation starting from day 2 and lasting till day 5 (Fig. 5G). CHOP^KD^ SKOV3 clones (Fig.S5A) were more resistant to IFN-γ, but not as much as the IDO1^KD^ cells, suggesting the possibility that there might be other factors involved in promoting TRP depletion induced cell death (Fig. 5H and Fig. S5B). These data demonstrate that the cell death caused by IFN-γ via TRP starvation is partially controlled by CHOP. Therefore, additional pathways downstream of IDO1 are involved in IFN-γ-mediated cell death that are yet to be described.

### IFN-γ synergizes with GCN2 inhibition in ovarian cancer

The ISR enables cells to adapt to different stress conditions and delays cell death. However, prolonged stress exposure ultimately leads to cell death (*44*). To test whether the ISR could be a possible target for HGSOC, we inhibited GCN2 in either SKOV3^WT^ or IDO1^KD^, using a specific inhibitor (GCN2-IN-6) (*37*) in combination with IFN-γ stimulation. Beginning at day 6, with a further increase by day 8, we observed increased cell death only in the WT cells treated with the combination treatment compared to the single treatments (Fig. 5I and S5C). In summary, our data suggests that IFN-γ induced TRP starvation in ovarian cancer cells sensitizes the malignant cells to GCN2 inhibitors. Thus, we conclude that GCN2 has two roles in the response to IDO1 expression and TRP depletion. First, the initial phase of GCN2-ISR activation is cell-protective. Later, upon prolonged IDO1 expression and chronic TRP depletion, the same pathway promotes cell death, in part by CHOP. All of these effects are reversible with IDO1i.

## Concluding remarks

The expectation from over two decades of research into how IDO1 suppresses T cell proliferation and function was that an IDO1i would reverse this pathway and ignite anti-tumor immunity. This rational guided the development and clinical evaluation of at least six IDO1i (*9, 10*). Yet, this approach overlooked a potential critical dimension: IDO1 direct, tumor intrinsic effects. We found that prolonged IDO1 expression promotes ovarian cancer cells death via p-GCN2/ATF4/CHOP axis, an effect reversible by exogenous TRP, inhibition of IFN-γ signaling, pharmacological or genetic IDO1 suppression, or modulation of the GCN2 ISR via CHOP depletion.

Our data support a model in which IFN-γ, likely from T cells, triggers IDO1 expression in the TME in a heterogeneous pattern. We hypothesize that within the tumor mass, IDO1^+^ cells will die while IDO1^-^ tumor cells, which are also present in high numbers in the TME, would survive. Administration of an IDO1i in this context could shift this balance, granting IDO1^+^ cells the same survival advantage as IDO1^-^ cells. Indeed, our results demonstrate that IDO1, amongst the hundreds of genes and proteins induced by IFN-γ, is responsible for its anti-tumor activity. Our results suggest that IDO1i result in the perverse outcome of promoting survival of tumor cells encountering IFN-γ.

Thus, IDO1i may have dual outcomes: restoring T cell activity but failing to overcome the survival advantage to tumor cells that escape the GCN2 ISR. Conversely, the TRP starvation-induced ISR can be targeted with GCN2-IN-6 to enhance tumor cell killing. A further unintended consequence of IDO1i use could be reduced neoantigen formation via ribosome skipping at TRP codons or TRP-to-Phe misincorporation (*45-47*). potentially dampening anti-tumor immunity. Thus, the framework for target validation of IDO1i, based solely on reversing T cell suppression, requires fundamental re-evaluation.

## Acknowledgements

We thank NGS facility at MPIB (RRID: SCR_025746) for the NGS sequencing service, the MS core facility at MPIB (RRID: SCR_025745) for metabolite measurements, the imaging core facility at MPIB (RRID: SCR_025739) for providing the CytoFLEX SRT Benchtop Cell Sorter. T.X., A.M., K.M. and C.D. were supported by the International Max Planck Research School for Molecular Life Sciences. E.L.L. was supported by the Medical Scientist Program of the Ludwig Maximilian University Munich (LMU) (MS-025). We acknowledge the use of Chat-GPT5 in helping us to review the writing of this manuscript. AI generated prompts were critically revised for accuracy.

## Author contributions

Conceptualization: P.J.M, M.M., T.X., A.M.

Methodology: T.X., A.M., X.Z., B.S. and E.L.L.

Formal analysis: T.X., A.M., X.Z., K.G.M., T.M.N. and E.L.L.

Investigation: T.X., A.M., L.K., B.S., L.S., C.D. and S.R.

Visualization: P.J.M., T.X., A.M., X.Z. and K.G.M.

Resources: E.L., E.O, Z.S.

Funding acquisition: P.J.M. and M.M.

Project administration: P.J.M. and M.M.

Supervision: P.J.M., M.M.

Writing – original draft: P.J.M., T.X. and A.M

Writing – review & editing: P.J.M., M.M., T.X. and A.M.

## Data and material availability

All spatial proteomics and RNA-seq data will be made publicly available.

**Supplementary materials**

Materials and Methods

Figs. S1 to S5

Tables S1 to S2

Additional references

