## Supplemental Material for "Spatial proteomics reveals mechanisms of cell-intrinsic tryptophan metabolism controlling ovarian cancer survival"

Peter J. Murray^1*^

*Corresponding author: Peter J. Murray

**The PDF file includes:**

Materials and Methods

Figs. S1 to S5

Tables S1 to S2

Additional references

Materials and Methods

***Human samples***

Treatment naive HGSOC specimens were obtained from four patients undergoing primary debulking surgery at the University of Chicago Medical Center. All patients provided written informed consent in accordance with institutional guidelines. Tumor diagnosis was confirmed by board-certified gynecologic pathologists. Fresh tissue was fixed in 10% neutral buffered formalin for 24 hours and paraffin-embedded following standard protocols. The study was approved by the Institutional Review Boards of the University of Chicago and the Max Planck Institute of Biochemistry.

***Deep visual proteomics***

FFPE tumor blocks were sectioned at 3 µm thickness onto laser microdissection-compatible PEN membrane slides (MicroDissect) treated with Vectabond (SP-1800-7, VectorLabs), according to the manufacturer’s instructions. Following deparaffinization, antigen retrieval (90°C, 40 min, pH 9) as described before (*1*), and blocking with 5% BSA in PBS for 1 hour, sections were immunostained overnight at 4°C with primary antibodies against IDO1 (1:100) and CD45, followed by secondary antibodies conjugated with alexa-647 (A21245, Invitrogen) and alexa-750 (A21037, Invitrogen) respectively (1:200 each, 1 hour RT) and incubation with alexa-555 directly conjugated EPCAM primary for two hours at 33°C and Hoechst nuclear counterstain. Slides were imaged at 20× magnification using widefield microscopy (Zeiss Axioscan 7).

Cell segmentation was performed using CELLPOSE 2.0 (*2, 3*)with EPCAM as primary and Hoechst as secondary channel, while downstream image analysis was performed on the BIAS software (*4*). Within EPCAM+ tumor cells, k-means clustering identified IDO1^hi^, IDO1^med^, and IDO1^lo^ populations based on IDO1 fluorescence intensity. CD45^+^ immune cells were excluded from analysis. Approximately 450 cell segments per group were laser micro-dissected (LMD7, Leica) and collected into 384-well plates. Samples were processed as previously published (*5*), specifically using automated liquid handling (Agilent Bravo), digesting with trypsin, and analyzing by LC-MS/MS on a Bruker timsTOF SCP mass spectrometer coupled to an Evosep One liquid chromatography system running a 20 samples/day (58-minute-long) gradient on an Aurora Elite CSI third generation column (IonOpticks, 15 cm x 75 μm, C18, 1.7 μm particle size, # AUR3-15075C18-CSI) at 50 ^o^C and 1300 V. The mass spectrometer was operated in diaPASEF mode with an optimal DIA isolation scheme generated by py_diAID (*6*), at an m/z range from 350 to 1200 and ion mobility from 0.7 to 1.3 Vs. cm-2 using 12 scans with two ion mobility windows per scan and a cycle time of 1.4 seconds. Raw MS data were analyzed using DIA-NN v1.8.1 (*7*) in library-free mode against the human proteome (UniProt: UP000005640_9606.fasta and UP000005640_9606_additional.fasta, accessed July 2021). Search parameters included: trypsin/p digestion with ≤1 missed cleavage, variable modifications (N-terminal Met excision, Cys carbamidomethylation, Met oxidation), peptide length 7-55 aa, precursor charge +1 to +4, m/z range 300-1800, and fragment m/z 200-1800. Match-between-runs and heuristic protein inference were enabled with 1% precursor FDR. Gene-level protein inference, single-pass neural network classification, robust LC quantification, and global normalization were applied. Statistical analyses and pathway enrichment were performed in R using limma for differential expression and WebGestalt for GSEA against Reactome databases.

***Spatial Proximity Analysis***

Following immunofluorescence staining and imaging, spatial relationships between tumor and immune cells were quantified using custom computational pipelines. Cell coordinates were extracted from the BIAS software output, representing the centroid positions of all segmented cells. For each tissue section, three distinct XML files containing cell coordinates were generated corresponding to IDO1^hi^, IDO1^med^, and IDO1^lo^ tumor cell populations, along with CD45^+^ immune cell positions.

Spatial analysis was performed using Python with the spatialdata and geopandas libraries. XML coordinate files were parsed and converted to spatial dataframes with consistent coordinate reference systems. For each tumor cell, the minimum Euclidean distance to the nearest CD45^+^ immune cell was calculated using the geopandas distance function. In total, distances were calculated for 17,158 IDO1^hi^, 219,520 IDO1^med^, and 2,166,381 IDO1^lo^ tumor cells relative to 90,784 CD45^+^ immune cells across all patient samples. These distances, reported in arbitrary units from the BIAS software coordinate system (*4*), served as a metric for tumor-immune cell proximity. Statistical comparisons of tumor-immune distances across IDO1 expression groups were performed in R. Two-sample t-tests were conducted for all pairwise comparisons between IDO1^hi^, IDO1^med^, and IDO1^lo^ groups within each patient sample (n=4 patients). The t-test was deemed appropriate given the large number of observations, which ensures robustness to deviations from normality through the central limit theorem. Results were considered statistically significant at p < 0.05. All distance measurements and statistical analyses were performed independently for each patient to account for inter-patient variability in tissue architecture and cell density.

***Mammalian cell culture***

SKOV3, OVCAR4, OVCAR5, OVCAR3, OVCAR8 and PA-1 cells were cultured in Dulbecco’s Modified Eagle Medium (DMEM) supplemented with 10% fetal bovine serum (FBS) and 1% penicillin-streptomycin (P/S) at 37°C in 5% CO2. TRP depleted DMEM was prepared starting from DMEM powder supplemented with 1% P/S, 3.7 g/L sodium bicarbonate, 4.5 g/L D-glucose, 0.1 g/L sodium pyruvate, 5% dialyzed FBS and all amino acids except for TRP. All cells were regularly tested negative for Mycoplasma contamination.

***Incucyte cell death assay***

The day before starting the assay, 12,500 SKOV3 cells were seeded in a 48-well plate. To start the assay, media was exchanged with fresh one containing Celltox Green (1:6000) and different treatments. All compounds were added only once at the same time unless stated differently. Cells viability was monitored by live phase-contrast microscopy using the IncuCyte S3 system with the 10X objective, pictures were taken every 24 hr up to 5 or 11 days. Cell death was quantified by counting the numbers of green fluorescence positive cells (CellTox Green positive) normalized to the confluence of each respective image.

3D assays were performed in 96- well ultra-low attachment plates, 5000 SKOV3 cells were seeded and centrifuged 300gx10min, on the same day all compounds and Celltox Green (1:6000) were added to the culture. Cells viability was monitored with the IncuCyte S3 with the 4X objective, pictures were taken every 24hr up to 5 days. Cell death was quantified by the green fluorescence intensity of each spheroid.

***CellTiterGlo assay***

SKOV3 (3500 cells/well), OVCAR4 (3000 cells/well), OVCAR5 (6500 cells/well), OVCAR8 (3000 cells/well) and PA-1 (6500 cells/well) cells were seeding in a 96-well plate. The day after, media was exchanged with one containing different compounds. The CelltiterGlo assay was performed after 4 days (PA-1), 5 days (SKOV3), 6 days (OVCAR4, OVCAR5 and OVCAR8), by adding 25 μL of the CellTiter-Glo® reagent into each well, after mixing cells were transferred to white opaque plates. Plates were shaken for 2 min at 180 rpm, rested for 10 min and luminescence was acquired using the Spark® plate reader (Tecan).

***Immunoblotting***

Cells were seeded on 6-well plate at different density depending on the length of the experiment (400,000 SKOV3 cells/well for 24h treatments, 200,000 SKOV3 cells/well for treatments longer than 4 days and 250,000 OVCAR4 cells/well for 30h treatments). At the end of the treatment period, cells were lysed with RIPA buffer containing 1% protease and phosphatase inhibitors. Cell lysates were separated on 4-15% polyacrylamide gel and transferred to 0.2 μm PVDF membranes. Afterwards, the blots were blocked with different blocking buffer diluted in TBS-T according to the primary antibody used (Table S2). Primary antibody incubation occurred overnight and the day after, blots were washed three times with TBS-T (10 min each wash), incubated with the respective secondary antibody coupled to HRP, followed by three washes and developed using the SuperSignal West Pico PLUS Chemiluminescent substrate.

***Plasmids***

SgRNA targeting different DNA loci were cloned into pSpCas9(BB)-2A-GFP vector using Bbs I restriction site and ligated with the T4 ligase reaction. The ligation product was transformed into competent cells. To generate the inducible hIDO1 and hIDO1F226A constructs, the respective cDNAs were amplified from in-house pcDNA3.1-hiDO1 and pcDNA3.1-hIDO1F226A and cloned into PB-TRE-dCas9-VPR instead of the dCas9 using the site between Nhe I and Age I. Meanwhile, mCherry-T2A-Flag-hypPB was synthesized with Twist Biosciences and cloned into pTwist backbone using the site between Not I and Xba I. The cloning procedure is the same as described before. All the constructs were confirmed by Sanger sequencing and whole plasmid sequencing.

***RNA isolation***

Cells were lysed using Trizol at -80 °C, followed by chloroform precipitation. The RNA fraction is precipitated with isopropanol and washed with 70% ethanol. Finally, the pellet was left to dry and resuspended in RNAse-free water.

***RNA-seq and analysis***

Bulk mRNA sequencing libraries were prepared with 500ng of total RNA of each sample using the NEBNext Ultra™ II Directional RNA Library Prep Kit for Illumina® (E7760, NEB) with NEBNext® Poly(A) mRNA Magnetic Isolation Module (E7490, NEB), according to standard manufacturer’s protocol. Total RNA and the final library quality controls were performed using Qubit™ Flex Fluorometer (Q33327, Thermo Fisher Scientific) and 4200 TapeStation System (G2991BA, Agilent) before and after library preparation. The libraries were sequenced on Illumina NovaSeq 6000 SP flow cell (2 x 60 bp) and demultiplexed by bcl2fastq Conversion Software (Illumina). After checking the quality of the samples (FastQC, v.0.12.1) the files were mapped to the human genome (Genome build GRCh38) downloaded from Ensembl using the star aligner version 2.7.10b. The mapped files were then quantified on a gene level based on the Ensembl annotations, using the HTseq tool69 in Python. Using the DESeq2 package70 (R 4.3.3, DESeq version 1.42.0), the count data was normalized by the size factor to estimate the effective library size. A filtering step of removing genes with less than 1 reads was used. This followed by the calculation of gene dispersion across all samples. The analysis of two different conditions against each other resulted in a list of differentially expressed genes for each comparison. Genes with a p-value of ≤ 0.01 and a log(2)Foldchange of > |1| were then considered to be differentially expressed for downstream analysis.

***Flow cytometry***

Cells were collected after trypsinization, washed twice with PBS and stained with 1 µl of APC-Annexin V and 2 µl of 7-AAD diluted in 50 µl/well binding buffer for 15 min at room temperature. Before flow cytometry analysis cells were diluted with 50 µl of binding buffer. Samples were acquired using the Cytek® Northern Lights™ with three lasers configuration (405 nm: 100 mW, 488 nm: 50 mW, 640 nm: 80 mW)

***Cell sorting***

Transfected cells were collected after trypsinization, washed twice with FACS buffer (PBS with 2% FBS), finally resuspended in FACS buffer and processed using the CytoFLEX SRT Benchtop Cell Sorter (Beckman Coulter). To generate knock-out cell lines GFP^+^ cells were sorted as single cells in 96-well plates. While, to generate stably Tet-on inducible hIDO1 cell lines, mCherry^+^ cells were enriched into 12-well plates for further selection.

***Targeted metabolites measurement***

Cells were plated at the density of 400,000 cells/well in 6 well plates, following 24 hr of IFN-γ stimulation (10 ng/µl) both supernatant and cell pellets were collected and snap frozen in dry ice. The supernatant was centrifuged 400g rcf x 5min to remove possible cell debris before freezing. Samples were stored in -80 °C until further processing.

Ice-cold extraction buffer (1 mL, ACN/MeOH/water 2:2:1 v/v) was added to 200 µl of supernatant. The samples were vortexed and ultrasonicated for 10 minutes, with 10 cycles of 30 seconds at high intensity followed by a 30-second pause between each cycle (Bioruptor, Diagenode). After centrifugation for 10 minutes at 12,000 rcf at 8°C, the supernatants were dried using a vacuum concentrator (SpeedVac). Cell pellets were extracted with the same protocol and extraction buffer with addition of 0.1M formic acid. Afterwards, 60 µl of a saturated ammonium bicarbonate solution (15%) was added. After centrifugation for 10 minutes at 12,000 rcf at 8°C, the supernatants were dried using a vacuum concentrator (SpeedVac).

Data were recorded using a QExactive HF mass spectrometer (Thermo Scientific) coupled with a Vanquish Flex HPLC system (Thermo Fisher Scientific). Vacuum-dried metabolite extracts from supernatants were dissolved in 160 μL of buffer A (0.1% formic acid), and 1 μL of the samples was injected. Vacuum-dried metabolite extracts from pellets were dissolved in 100 μL of buffer A (0.1% formic acid), and 3 μL of the samples was injected. For the detection of Kyn and Trp, metabolite extracts were separated on a Kinetex F5 column (2.1 x 100mm, 2.6 μm, Phenomenex) at a flow rate of 200 µL/min. Buffer B (100% ACN, 0.1% formic acid) was maintained at 0% for 2 minutes, then increased to 95% within 12 minutes, and held at 95% for 2 min. The column was then re-equilibrated to 0% B for 4 minutes. For the detection of NAD and NADH, metabolite extracts were separated on a Luna NH2 column (2.1 x 100 mm, 3.0 μm, Phenomenex) at a flow rate of 300 µL/min. The gradient started with 95% of buffer B (100% ACN) and 5% of buffer A (95% ACN, 20 mM ammonium acetate, 20 mM ammonium hydroxide). Percentage of buffer A was increased to 100% within 6 minutes, was maintained at 100% for 9 minutes, then decreased to 5% within 1 minute, and held at 5% for 7 minutes.

The mass spectrometer operated in positive mode for the detection of Kyn and Trp and in negative mode for the detection of NAD and NADH. Data-dependent MS1 scans were recorded from 75 to 1000 m/z at a resolution of 120,000. Conditions for the HESI source were as followed: Sheath gas (N2) flow rate was set to 47 (arbitrary units), auxiliary gas flow rate was set to 10 (arbitrary units), and sweep gas flow rate was set to 2 (arbitrary units). The spray voltage was maintained at 3.20 kV, and the capillary temperature was set to 250 °C and the temperature of the auxiliary gas heater was set to 380°C. Up to 5 of the top precursors were selected and fragmented using higher energy collisional dissociation (stepped-HCD with normalized collision energies of 20, 40, and 60). The MS2 spectra were recorded at a resolution of 30,000. The AGC target for MS1 and MS2 scans was set to 3E6 and 1E5, respectively, within a maximum injection time of 200 ms for MS and 50 ms for MS2 scans. Peak areas corresponding to Kyn, Trp, NAD and NADH were identified based on MS1 high-resolution mass, characteristic MS2 fragments, and retention time (previously identified using unlabeled standards) using the software "Skyline," version 23.1.0.455 (MacCoss lab, University of Washington). Concentrations were calculated using previously recorded calibration curves of un-labelled standards.

***scRNA-seq data analysis***

Single-cell RNA sequencing (scRNA-seq) data were obtained from a publicly available ovarian cancer dataset, which can be downloaded from the original publication (DOI: 10.1038/s41586-022-05496-1). Data processing followed the computational pipeline described in the source article: reads were aligned, filtered, and quantified using CellRanger (v3.1.0). Quality control steps included retaining cells with at least 500 expressed genes, over 1,000 UMI counts, and less than 25% mitochondrial gene expression. Doublets were filtered using Scrublet (v0.2.1) with a threshold of 0.25, and cell cycle phases were assigned using Seurat’s CellCycleScoring function. Sample matrices were merged by patient and renormalized using default Seurat functions. Major cell types were identified with CellAssign (v0.99.2) based on curated marker genes for nine ovarian cancer-associated cell types, as in the original study. Cells lacking expression of all marker genes were excluded. Dimensionality reduction was performed using PCA, and UMAP embeddings were generated from the first 50 principal components. Batch correction was conducted using the Harmony package to account for patient-specific effects. To investigate the expression patterns of IDO1 and IFNG mRNAs across different cell types, we calculated both the average expression and the percentage of expressing cells for each gene within each annotated cell type. IDO1 and IFNG expression were mapped onto the UMAP embedding to illustrate their distribution within the cellular landscape. Unlike conventional feature plots, these density-based visualizations were generated using the Nebulosa R package. Pathway activity scores for the ISR and ISGs were computed for each cell using the AddModuleScore function, based on curated gene sets for each pathway (detailed table in supplementary material). To assess the relationship between IDO1 expression and pathway activity, tumor cells were extracted and divided into two groups according to their IDO1 expression status: cells with detectable IDO1 transcripts were assigned to the "high" group, while those without detectable expression were assigned to the "low" group. The activity scores of the ISR and ISGs pathways were then compared between these two groups. Differences in pathway scores were visualized using violin and box plots, and statistical significance was assessed using the Wilcoxon rank-sum test. All analyses were performed using Python and R, leveraging Scanpy, Seurat, and related packages for scRNA-seq data processing, visualization, and statistical testing.

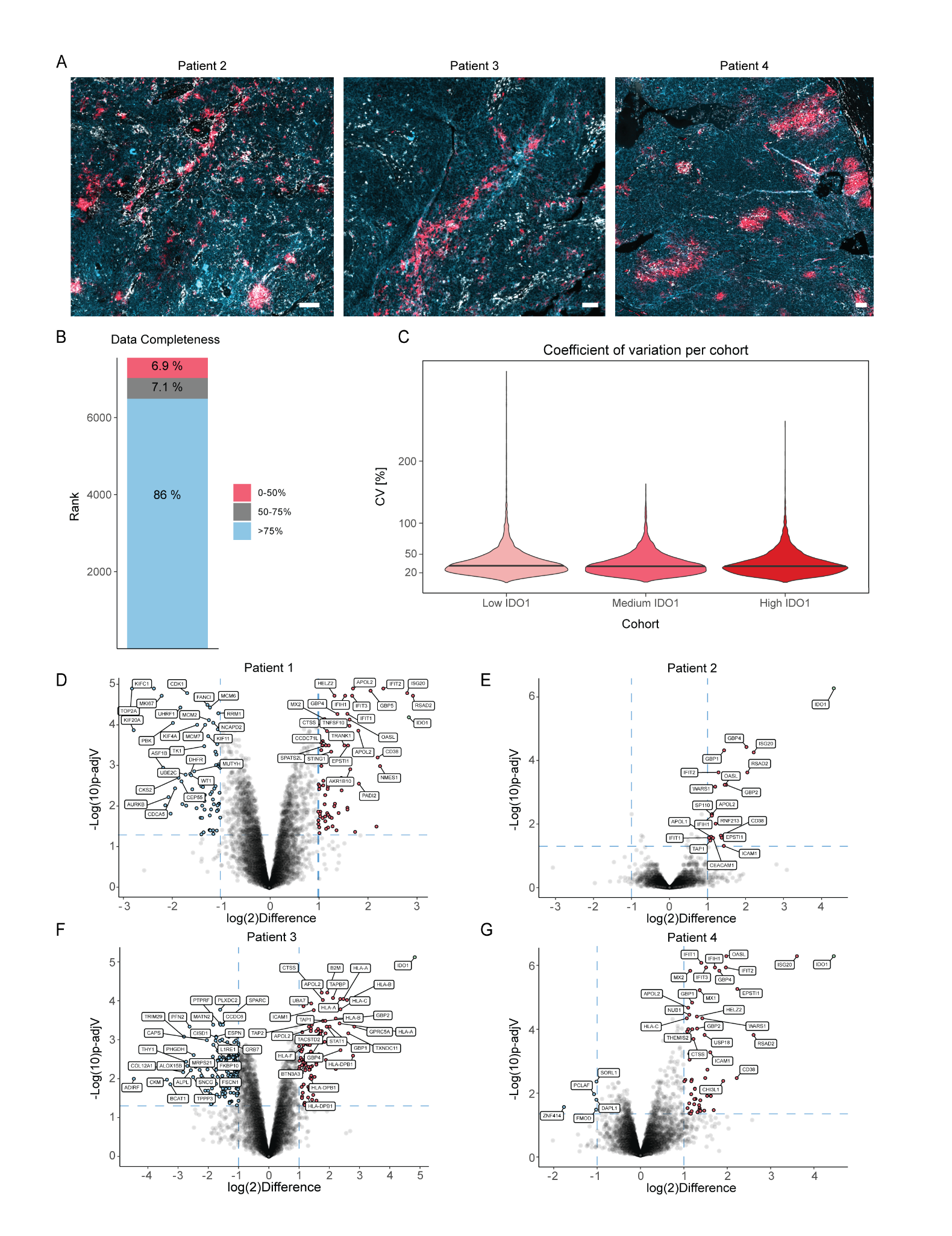

**Fig. S1.** **IDO1 heterogeneity and proteomics data quality across HGSOC patients**. (**A**) Representative immunofluorescence images showing heterogeneous IDO1 expression patterns across patients 2, 3, and 4. Scale bars = 100 μm. Cyan: EPCAM, red: IDO1, white: CD45. (**B**) Data completeness showing the percentage distribution of proteins with different coverage across all samples. The percentage of proteins identified in >75% (blue), 50-75% (gray), and <50% (red) of all samples, respectively. (**C**) Coefficient of variation (CV) for proteins across IDO1 expression cohorts. CV distribution for proteins with valid values across IDO1 expression groups. Violin plots display the density of CV values. (**D** to **G**) Volcano plots showing differentially expressed proteins between IDO1^hi^ (right) and IDO1^lo^ (left) tumor cells for each patient (P1-P4). Gray dots represent proteins with non-significant changes, thresholds for significance are log(2)Difference < -1 or >1 and -Log(10)p-adjV > 1.3, red = upregulated in IDO^hi^ and blue = downregulated in IDO1^hi^. The top 25 up- and downregulated proteins were annotated.

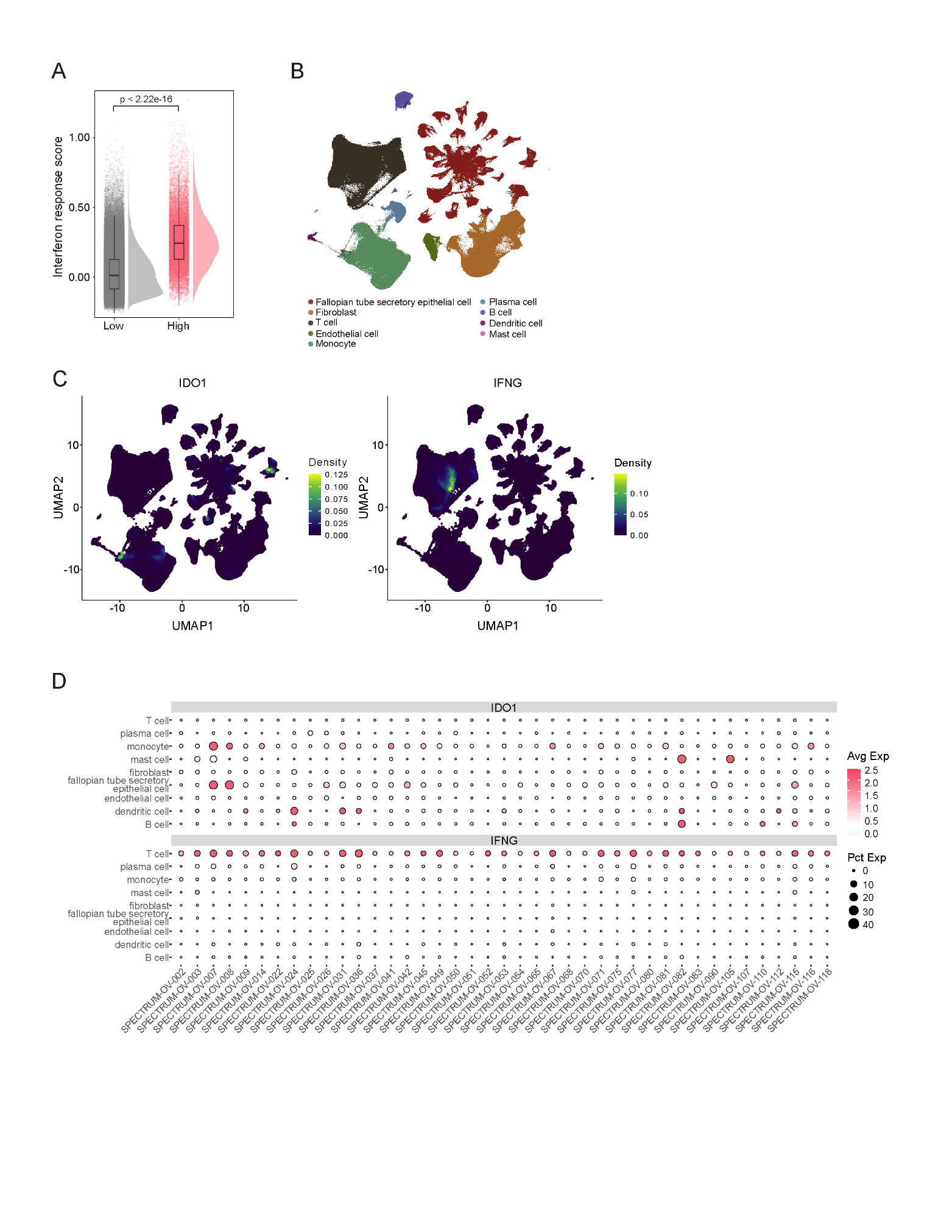

Fig. S2. IDO1 and IFN-γ expression across different HGSOC patient tissue. (A) Density curves and boxplots show interferon response score per tumor cell in the IDO1^lo^ and IDO1^hi^ expressing group. Significant p-value was calculated with the Wilcoxon rank-sum test. (B) ScRNA-seq analysis of HGSOC patient data, UMAP dimensional reduction colored by cell types (C) scRNA-seq analysis showing IDO1 enrichment in tumor epithelial cells and monocytes, while IFNG is enriched in T cells. (D) Dotplot shows scaled and averaged expression levels of IDO1 and IFNG per cell type in each patient. Dot size reflects the fraction (%) of expressing cells; color scaled means expression level.

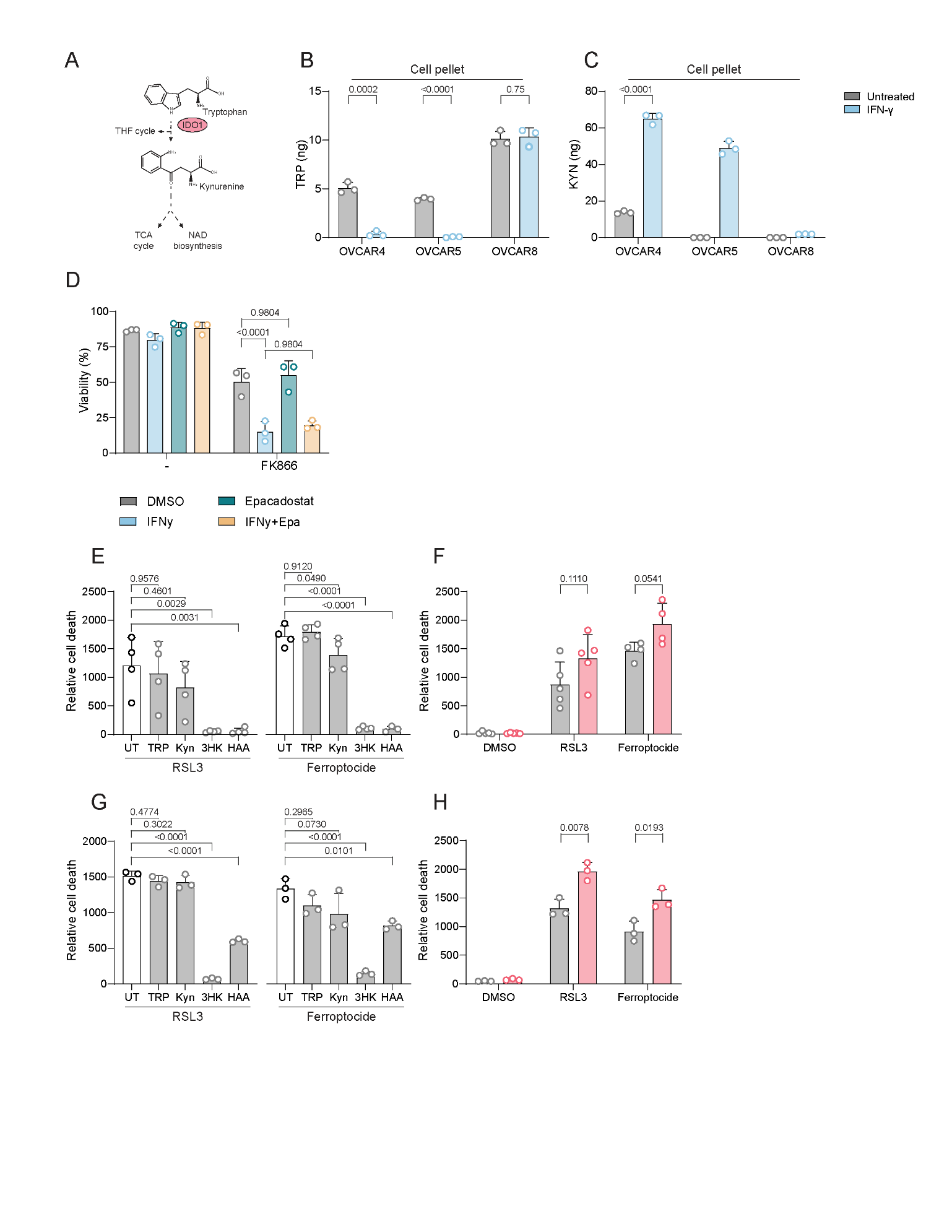

Fig. S3. Exclusion of IDO1 dependent NAD de novo synthesis and ferroptosis protection. (A) Schematic representation of the KP pathways fueling to THF cycle (1-carbon metabolism), TCA cycle and NAD biosynthesis. (B to C) TRP (B) and KYN (C) measured by mass spectrometry in the cell pellet of OVCAR4, OVCAR5 and OVCAR8 stimulated with 10 ng/ml of IFN-γ for 24hr. (D) Percentage viability calculated as annexin V^-^ and 7-AAD^-^ of SKOV3 cells treated for 5 days with either vehicle (DMSO), 10ng/ml IFN-γ, 10 µM epacadostat or both in combination with 1 µM FK866. (E and G) Quantification of live cell imaging data as relative cell death (Count of CellTox Green positive cells/Confluence) of SKOV3 (E) treated with either 1 µM RSL3 or 10 µM ferroptocide; while OVCAR4 (G) were treated with either 0.25 µM RSL3 or 2.5 µM ferroptocide. At the same time treated cells were cultured media containing different KP metabolites (200 µM KYN for ferroptocide treated cells and 100 µM KYN for RSL3 treated cells, 25 µM 3-HK and 25 µM 3-HAA) and 200 µM TRP was used as control. (F and H) Relative cell death of WT versus IDO1 depleted SKOV3 (F) or OVCAR4 (I) treated with either RSL3 or ferroptocide (concentrations as above). All data are shown as means ± SD of at least three independent experiments. Unpaired t-test was used in (B), (C), (F) and (H). Two-ways ANOVA complemented with a Tukey’s multiple comparison test in (D). One-way ANOVA complemented with a Dunnett’s multiple comparison test in (E) and (G). One data point in (H) was excluded in the ferroptocide treated cells as defined as outlier by the Grubbs test (alpha = 0.05).

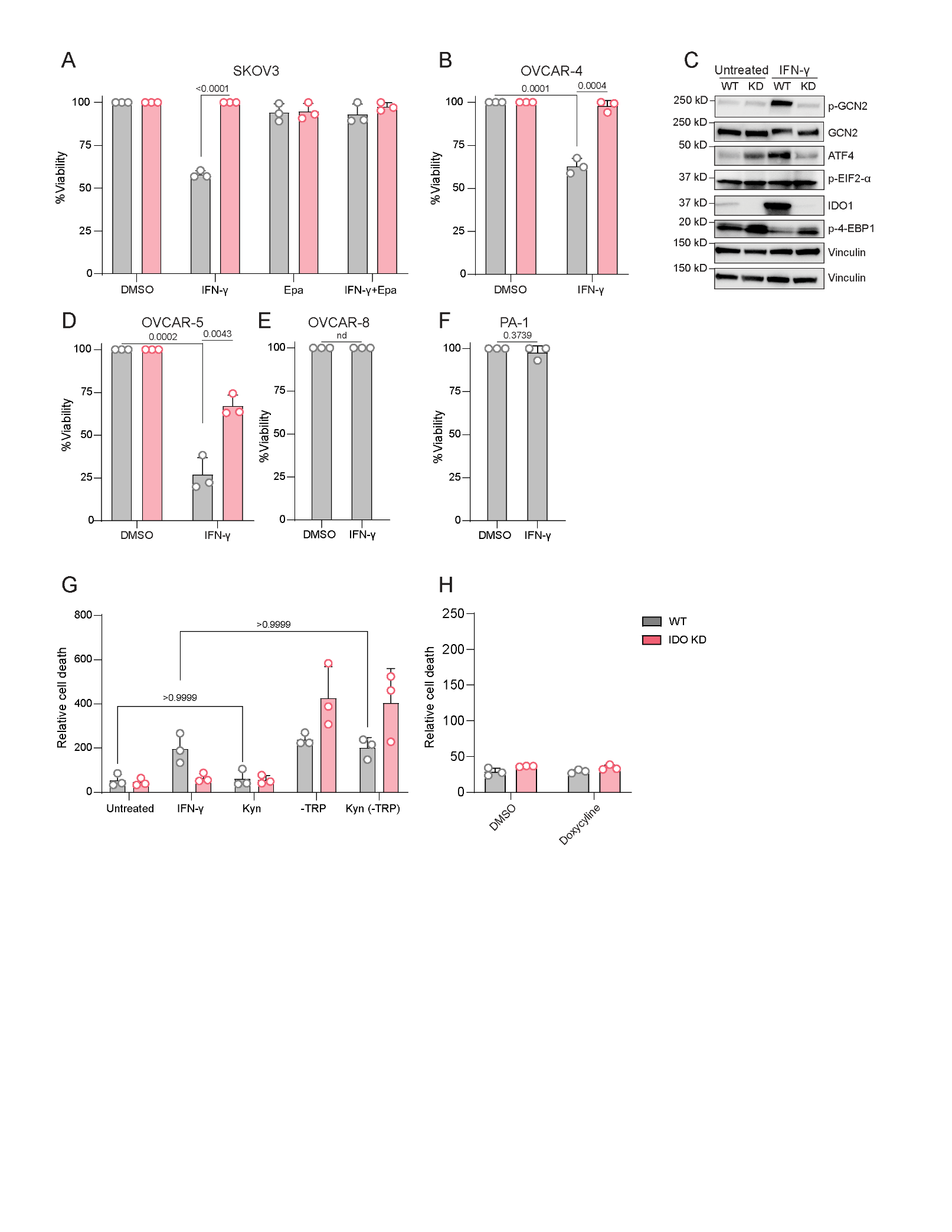

**Fig. S4. IDO1 dependent IFN-γ killing and ISR activation across multiple ovarian cancer cell lines.** (**A**) Cell-titerGlo (CTG) assay of SKOV3^WT^ or IDO1^KD^ treated for 5 days with either vehicle (DMSO), 10 ng/ml IFN-γ, 10 µM epacadostat (Epa) or both, plot depicts viability in percentage relative to the untreated control. (**B** and **D**) Percentage viability from CTG assay of WT versus IDO1^KD^ OVCAR4 (B) or OVCAR5 (D) treated for 6 days with 10ng/ml and 50 ng/ml of IFN-γ, respectively. (**C**) Immunoblot analysis of the ISR pathway in OVCAR4^WT^ versus OVCAR4 IDO^KD^ untreated or stimulated with 10 ng/ml IFN-γ for 30 hr (n=3). (**E** and **F**) Percentage viability from CTG assay of OVCAR8 (E) treated for 6 days with 50 ng/ml of IFN-γ or PA-1 (F) treated for 6 or 4 days with 50 ng/ml of IFN-γ, respectively. (**G**) Relative cell death of SKOV3^WT^ versus SKOV3 IDO1^KD^ left untreated, or treated with either 10 ng/ml IFN-γ or 200 µM KYN, or cultured in DMEM without TRP +/- 200 µM KYN. This plot is linked to Figure 5C in the main manuscript. (**H**) Relative cell death of SKOV3^WT^ versus SKOV3 IDO1^KD^ treated with 20 ng/ml for 5 days of doxycycline. These are the controls for Figure 5F in the main manuscript, the y-scale is matching Figure 5F. All data are shown as means ± SD of at least three independent experiments. Unpaired t-test was used in (A), (B), (D), (E) and (F). Two-ways ANOVA complemented with a Tukey’s multiple comparison test in (G).

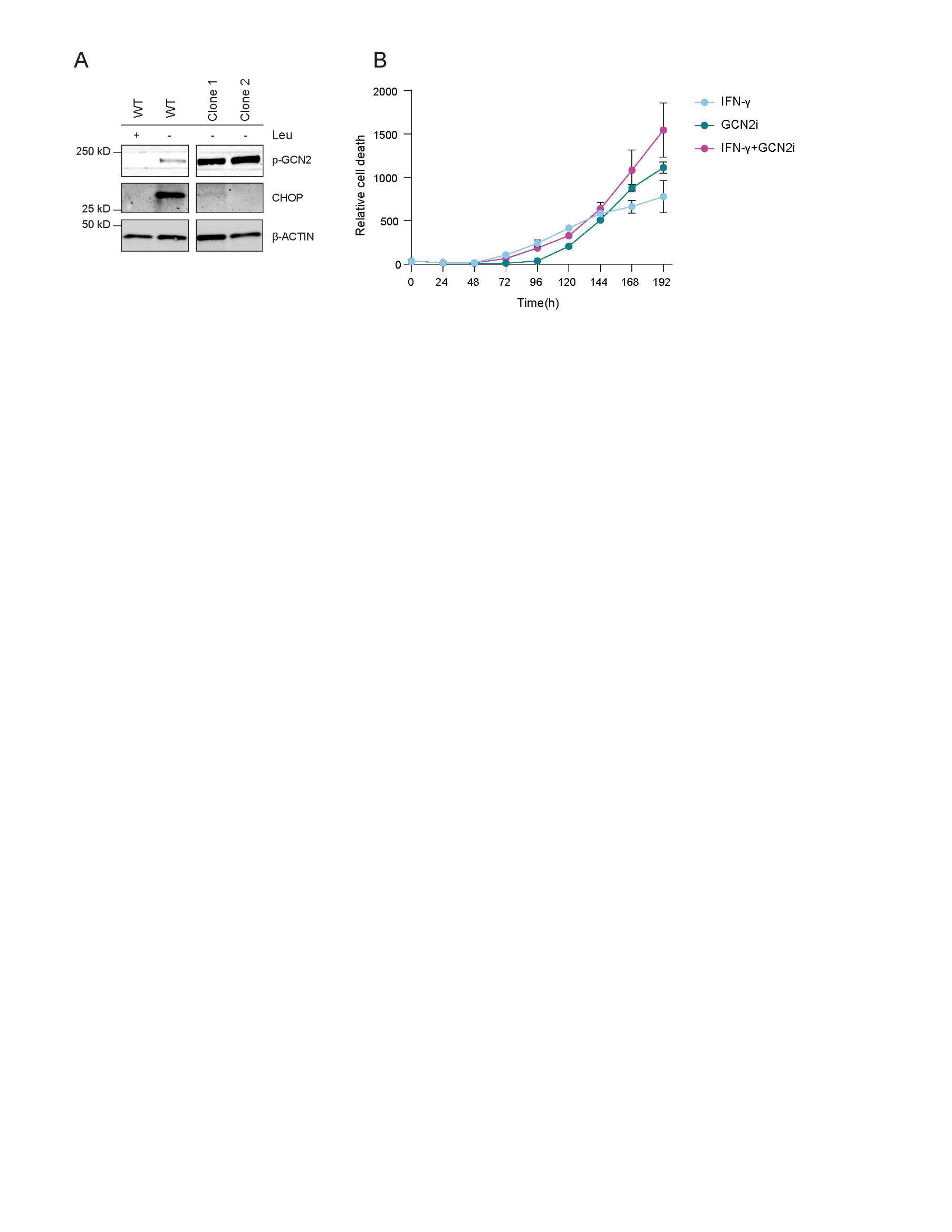

**Fig. S5. IDO1 dependent IFN-γ killing is partially regulated by CHOP and GCN2 inhibition synergize with IFN-γ.** (**A**) Immunoblot analysis of CHOP in WT or CHOP KO clones, to visualize CHOP expression, cells were leucine starved for 16hr. (**B**) Time-course relative cell death of SKOV3 treated with either 10 ng/ml IFN-γ, 1 µM GCN2-IN-6 (GCN2i) or both. Data shown as means ± SD of three technical replicate, this is a representative plot of four independent experiments.

| **Targets comprising amino acid starvation score** | | | |
| --- | --- | --- | --- |
| TRIB3 | GADD45A | FAU | RRAGB |
| ASNS | STC2 | FLCN | RRAGC |
| WARS | ACOT1 | FNIP1 | RRAGD |
| HERPUD1 | BCAT1 | FNIP2 | SEC13 |
| CHAC1 | IL1RL1 | GCN1 | SEH1L |
| SLC6A9 | EPRS | IMPACT | SESN1 |
| ATF3 | VLDLR | ITFG2 | SESN2 |
| ATF5 | PPP1R15A | KICS2 | SH3BP4 |
| ERO1A | ALDH1L2 | KPTN | SLC38A9 |
| CTH | ATF4 | LAMTOR1 | SZT2 |
| MTHFD2 | ATF2 | LAMTOR2 | TCIRG1 |
| SHMT2 | BMT2 | LAMTOR3 | UBA52 |
| PSAT1 | CASTOR1 | LAMTOR4 | WDR24 |
| GARS | CASTOR2 | LAMTOR5 | WDR59 |
| NARS | CEBPB | MIOS |  |
| PSPH | CEBPG | MLST8 |  |
| SLC3A2 | DDIT3 | MTOR |  |
| CBS | DEPDC5 | NPRL2 |  |
| CARS | EIF2AK4 | NPRL3 |  |
| GPT2 | EIF2S1 | RHEB |  |
|  | EIF2S2 | RPTOR |  |
|  | EIF2S3 | RRAGA |  |

Table S1. Starvation gene list

| **Antibody** | **Provider** | **Cat. Number** | **Blocking buffer** | **Dilution** |
| --- | --- | --- | --- | --- |
| Vinculin (E1E9V) XP® | CST | 13901S | TBS-T 3% milk | 1:1000 |
| GRB2 | BD bioscience | 610112 | TBS-T 3% milk | 1:1000 |
| IDO (D5J4E) | CST | 86630S | TBS-T 3% milk | 1:1000 |
| ATF-4 (B-3) | Santa Cruz | sc-390063 | TBS-T 5% BSA | 1:500 |
| GCN2 | CST | 3302S | TBS-T 3% milk | 1:1000 |
| p-GCN2 (T899) | Abcam | ab75836 | TBS-T 3% milk | 1:1000 |
| P-eIF2alpha (S51) (119A11) | CST | 3597S | TBS-T 1% BSA | 1:1000 |
| CHOP (L63F7) | CST | 2895S | TBS-T 3% BSA | 1:1000 |
| β-Actin (8H10D10) | CST | 3700S | TBS-T 3% milk | 1:1000 |
| eIF2alpha (D7D3) XP(R) | CST | 5324S | TBS-T 3% milk | 1:1000 |
| P-4E-BP1 (T37/46) (236B4) | CST | 2855S | TBS-T 3% milk | 1:1000 |
| 4E-BP1 (53H11) | CST | 9644S | TBS-T 3% milk | 1:1000 |
| CD45 | Dako | M0701 |  | 1:100 |
| Alexa Fluor® 555 Anti-EPCAM | Abcam | Ab275122 |  | 1:50 |

Table S2. Antibody list
